# Membrane Kymograph Generator: A cross-platform GUI software for automated generation and analysis of kymographs along dynamic cell boundaries

**DOI:** 10.64898/2026.02.11.705379

**Authors:** Tatsat Banerjee, Bedri Abubaker-Sharif, Peter N. Devreotes, Pablo A. Iglesias

## Abstract

**Summary:** The plasma membrane and accompanying cortex serve as one of the major hubs of the signal transduction and cytoskeletal activities that collectively regulate numerous cell physiological processes such as migration, polarity, macropinocytosis, phagocytosis, cytokinesis, etc. Yet, dynamically tracking membrane-cortex associated protein or lipid kinetics over time from live-cell image series remains a challenging task, primarily due to the difficulty of accurately extracting and aligning the cell boundary between consecutive frames, as the cell continuously deforms and moves. Here, we present *Membrane Kymograph Generator*, a cross-platform software that accepts multichannel time-lapse live-cell fluorescent imaging datasets as input and automates the cumbersome, heuristic process of boundary tracking, inter-frame alignment, and intensity sampling along the boundary. The software implements a rotational offset minimization algorithm that circularly aligns boundaries across consecutive frames by exhaustively searching for the optimal angular shift that minimizes point-to-point distances, while handling variations in boundary point counts due to cell shape changes. The software outputs kymographs that represent spatiotemporal dynamics of different membrane-associated proteins or biosensors, allows users to fine-tune visualization parameters through an interactive interface, and provides built-in correlation analysis tools for multi-channel datasets. Furthermore, the software allows advanced programmatic usage for batch processing and further analysis via a native API. Our validation tests demonstrated that the *Membrane Kymograph Generator* can be used to accurately track, visualize, and quantitate the spatial kinetics of a wide array of membrane proteins and lipid biosensors over extended time periods, in a variety of cell types, including *Dictyostelium* amoeba, human neutrophils, mouse macrophages, and different mammalian cancer cells. The GUI-based software is user-friendly, does not require any technical expertise from users, and significantly reduces the manual effort and time required for kymograph generation and downstream analysis, while ensuring high accuracy and reproducibility.

**Availability and Implementation:** Membrane Kymograph Generator is a free and open-source software, licensed under GNU General Public License 3.0 or later. This software is cross-platform: It can be graphically installed on both x86-64 and AArch64/ARM64 computers, running either Windows, macOS, or any standard Linux distribution. The software is distributed as single installer files (and portable executables) targeting specific hardware architectures and operating systems, and hence, it can be installed natively without any dependency resolution. The source code, detailed documentation, specific installers, portable binaries, and test data are freely available at https://github.com/tatsatb/membrane-kymograph-generator. Additionally, since the software is written in Python, it can also be installed inside any Python environment using PIP package manager (package ID: https://pypi.org/project/membrane-kymograph) and can also be interacted via a built-in Python API.

## 1 INTRODUCTION

The plasma membrane and associated cortex are dynamic structures that often undergo rapid remodeling in response to environmental cues or internal stochastic fluctuations. These asymmetric, transient remodeling events, stemming from the coordinated activities of various lipid and membrane protein components, collectively define the spatiotemporal dimensions of different signal transduction and cytoskeletal activities, and thereby regulate a plethora of cell physiological responses (Ridley et al., 2003; Swaney et al., 2010; Banerjee et al., 2025). With the recent advances in live-cell fluorescence microscopy approaches, genetically-encoded biosensors, and synthetic biology/optogenetic tools, it became easier to closely monitor and/or perturb specific membrane/cortex components with high spatiotemporal resolutions, inside live cells, in real-time. Despite the wealth of imaging data generated, quantitative analysis of membrane dynamics remains challenging. The curved, often complex geometry of cell membranes makes it difficult to apply standard image analysis methods designed for Euclidean coordinates. Furthermore, typical live-cell microscopy images report both morphological changes in the membrane as well as alterations in the dynamics of multiple membrane proteins or lipids — making it difficult to decouple these effects.

Kymographs, the two-dimensional space-time plots that can represent spatial position and fluorescence intensity over time, have widely emerged as a powerful visualization tool in cell biology, systems biology, and biophysics research domains. While kymographs (and downstream quantitative analyses) helped to dissect the intricate spatiotemporal dynamics of various cellular components, generating accurate kymographs along cell membrane/cortex from time-lapse live-cell imaging datasets remained a non-trivial task, especially as cell membrane undergoes large-scale deformations and remodelling during dynamic cell physiological processes such as migration, polarity, macropinocytosis, phagocytosis, cytokinesis, vesicular trafficking, etc. Due to significant progress in deep learning and associated approaches in the past decade, many important software tools, such as ilastik (Berg et al., 2019), CellPose (Stringer et al., 2021), CellTracker (Hu et al., 2021), Celldetective (Torro et al., 2025), DeepCell (Greenwald et al., 2022), and CellProfiler (Stirling et al., 2021; Kamentsky et al., 2011) have emerged to facilitate segmentation of whole-cell and organelles from live-cell imaging datasets. Although these tools can often accurately segment the whole-cell and/or organelle boundaries, they do not provide any functionality to automatedly track and align boundaries which often assume substantially different sizes and geometries between subsequent frames. Furthermore, existing kymograph plugins of Fiji/ImageJ (Schindelin et al., 2012), such as *KymographBuilder* (Mary et al., 2016) and *Multi Kymograph* (Rietdorf and Seitz, 2017) cannot generate kymographs along dynamic cell boundaries, as they are designed to work with static, user-defined straight or curved lines. While membrane kymograph has become an incredibly useful tool to visualize and quantitate dynamic cell physiological processes in recent years, researchers had either largely relied on manual or semi-automated approaches to extract and align dynamic cell boundaries (which are time-consuming, prone to user bias, may not accurately capture the complex dynamics, and cannot take advantage of modern segmentation tools), or focused on analyzing a straight-line segment across the membrane (which essentially represents incomplete 1D picture), or developed custom codes for specific applications (which are not user-friendly, generalizable, and often inaccessible to a significant fraction of experimental biologists) (Iwamoto et al., 2025; Arai et al., 2010; Deng et al., 2025; Gong et al., 2024; Banerjee et al., 2023; Lee et al., 2020; Bosgraaf and Van Haastert, 2010; Tyson et al., 2010; Tong et al., 2023; Kuhn et al., 2025; Baniukiewicz et al., 2018; Chua et al., 2024; Lin et al., 2024; Pal et al., 2023; van Haastert et al., 2017; Ghabache et al., 2021; Miao et al., 2017; Bosgraaf et al., 2009; Cheng and Mullins, 2020; Banerjee et al., 2022; Barry et al., 2015; Xiong and Iglesias, 2010). Additionally, some of these tools require user registrations and were not released under an approved free/libre license (Free Software Foundation, 2025). To the best of our knowledge, no free and open source software exists to fully automate the process of generating membrane kymographs from live-cell imaging datasets while accounting for the dynamic nature of cell boundaries.

Here, we present *Membrane Kymograph Generator*, a GNU GPL v3 licensed, cross-platform, end-to-end GUI standalone software that automates this entire process of generating and analyzing kymographs along dynamic boundaries from multichannel time-lapse live-cell fluorescent images. This software automates the extraction of the cell boundary, tracks and aligns the boundaries across frames using a rotational offset minimization algorithm, samples fluorescence intensities along the boundary as per user provided parameters, applies lowess smoothing and proper normalization, exports outputs in various accessible formats (for further analysis using other open-source/custom tools), generates publication-quality kymographs for each channel, and provides built-in correlation analysis tools for multi-channel datasets. In addition, we also implemented a simple Python-based API for programmatic access, batch processing, and further downstream analysis. Importantly, unlike previous tools that focus on specific cellular structures, or require knowledge in MATLAB/Python/Julia to perform extensive parameter tuning, *Membrane Kymograph Generator* provides an intuitive core interface suitable for researchers with any levels of computational or technical expertise.

## 2 SOFTWARE ARCHITECTURE AND RESULTS

*Membrane Kymograph Generator* is implemented as a Python package with additional components in bash and inno setup scripts. In addition to standard Python libraries, this software uses: `NumPy`, `SciPy`, and `Pandas` for numerical operations and data handling; `scikit-image`, `OpenCV`, `Shapely`, and `Pillow` for image processing tasks; `Matplotlib` and `Seaborn` for visualization; `joblib` for parallel processing; and `ttkbootstrap` (an enhanced `tkinter` framework) for building the modern graphical user interface. The modular architecture allows researchers to use all the tools via the integrated GUI application as well as programmatically via a built-in Python API for batch processing and advanced usages. A simple representation of the GUI workflow of *Membrane Kymograph Generator* is illustrated in Figure 1.

**Figure 1.**
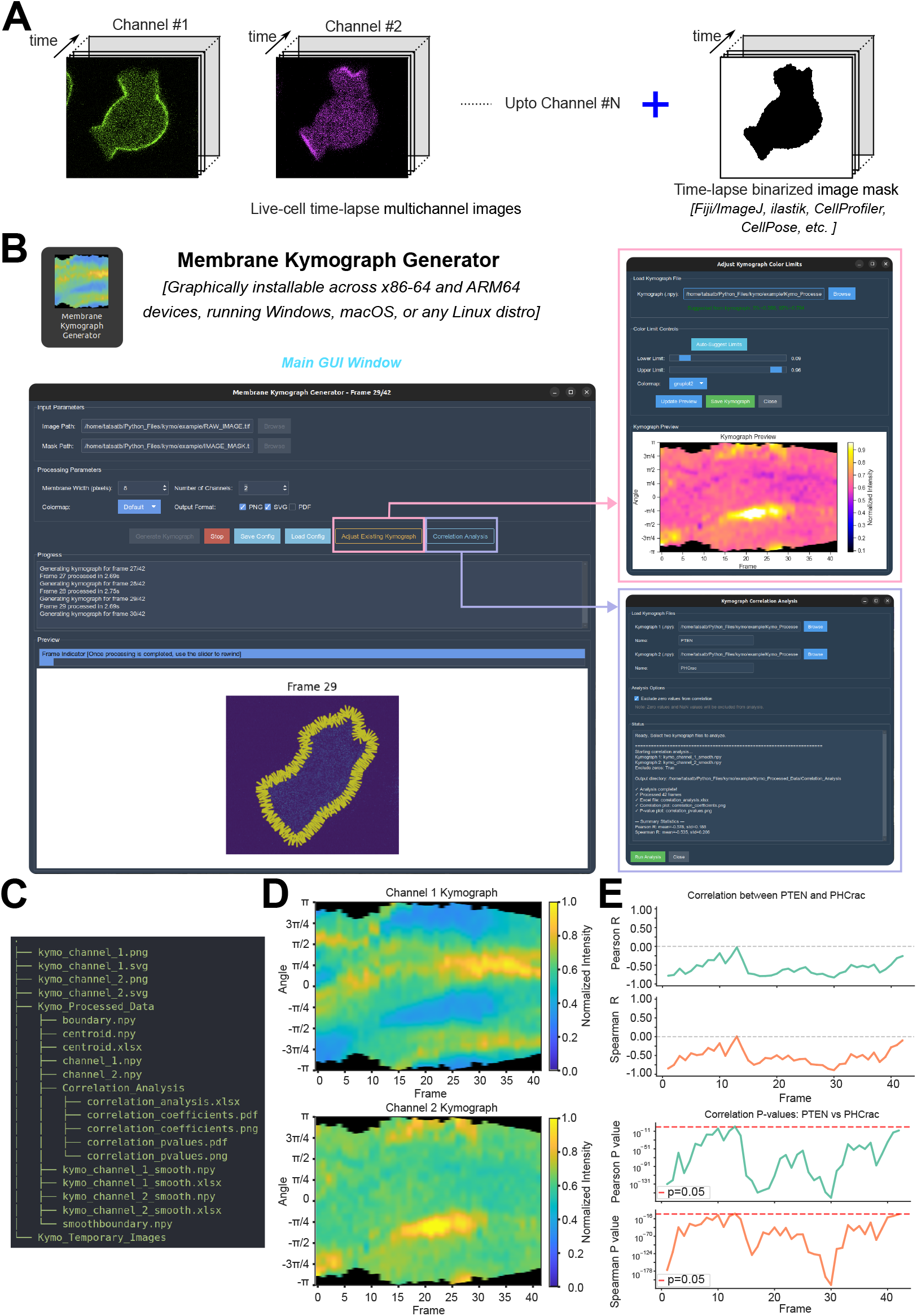
Overview of the default GUI workflow for Membrane Kymograph Generator. **(A)** Input files for the Membrane Kymograph Generator include a multichannel time-lapse live-cell imaging dataset and a binarized mask of the whole cell. Note that the binarized mask can be generated using various standard tools such as Fiji/ImageJ, ilastik, CellProfiler, CellPose, etc., or using custom code involving thresholding, morphological operations, or machine learning-based approaches. **(B)** Graphical User Interface of Membrane Kymograph Generator. The main interface is shown in the left whereas auxiliary windows are shown in the right. The main interface allows users to upload their input files, configure analysis parameters, and visualize the progress and error messages (if any) in real-time, and finally visualize the generated kymographs. The bottommost panel here shows the tracked boundaries and intensity sampling domain. The top right auxiliary window allows the users to adjust the visualization settings, such as colormap and intensity scaling. The bottom right auxiliary window facilitates the downstream correlation analysis across different kymographs. **(C)** List of key output files generated by the Membrane Kymograph Generator. The temporary images inside the *Kymo_Temporary_Images* folder are hidden from the tree view here. **(D)** Representative output kymographs generated by Membrane Kymograph Generator from a two-channel live-cell images over 42 frames. Evidently, each column corresponds to a specific time point, and each row corresponds to a specific position along the cell boundary. The intensity values in the kymographs indicate the fluorescence intensity of the respective proteins at each position and time point. The used colormap is shown on the right. **(E)** Representative output correlation analysis plots generated by the “Kymograph correlation Analysis” module of Membrane Kymograph Generator. The top two plots show the Pearson’s and Spearman correlation coefficient (R) values between two kymographs over time, indicating the degree of correlation between the two proteins’ dynamics along the membrane boundary. The bottom two plots show respective p-values associated with the correlation coefficients, indicating the statistical significance of the observed correlations.

### 2.1 Installation and updates

The *Membrane Kymograph Generator* can be installed on computers with both x86-64 and ARM64 architectures, running either Windows, macOS, or any standard Linux distribution. The software is primarily distributed as standalone installable packages (.exe for Windows,.dmg for macOS, and.appimage for Linux). Additionally, portable binary executables and/or tarballs for each supported platform are also provided. The codebase and installation had been thoroughly tested on Windows 10/11, Ubuntu 22.04 LTS/24.04 LTS, and macOS 15. For programmatic access and terminal-based workflows, the software can also be installed inside any standard Python environment (for e.g. conda, venv, pyenv/poetry, etc.) using PIP package manager from the Python Package Index (PyPI) repository (package ID: *membrane-kymograph*) or directly building from the source code. We have also implemented a *GitHub Actions* based *continuous integration and continuous deployment* (CI/CD) pipeline to automatically build, test, and deploy the new software versions upon every tagged commit to the main branch of the GitHub repository. Before installing a new version, users may prefer to uninstall any previous versions, simply via control panel (Windows), dragging to trash (macOS), or deleting the AppImage file or using provided uninstallation shell script (Linux).

### 2.2 Graphical User Interface overview and input files

The primary way of interacting with the software is via the graphical user interface (GUI) built using `ttkbootstrap/tkinter`, which provides an intuitive and modern interface for general users (Figure 1B-E). The main interface features organized input parameter panels where users can select their multichannel time-lapse image file and corresponding binarized mask file via OS native file browser dialogs. The processing parameters panel enables users to configure key analysis settings, including the membrane width for perpendicular intensity sampling (default: 8 pixels, adjustable), the number of fluorescent channels to process (1-10 channels), the desired colormap for visualization (offering multiple perceptually uniform and diverging colormaps including the default modified Parula, as well as viridis, plasma, inferno, gnuplot, and others), and the output file formats (PNG, SVG, and/or PDF). A progress panel provides real-time feedback during processing, displaying status messages and error diagnostics to keep users informed throughout the analysis workflow. The bottommost preview panel displays the tracked cell boundaries overlaid with perpendicular sampling lines on the original fluorescence images, allowing users to visually verify the accuracy of boundary detection, alignment, and intensity sampling across frames, as each frame is processed. This preview panel also shows the generated kymographs after the end of processing. An integrated frame slider enables users to navigate through the time series and inspect tracking quality. In addition to the main interface, auxiliary dialog windows provide advanced functionality: the “Adjust Existing Kymograph” dialog allows users to interactively fine-tune intensity scaling and colormap settings for previously generated kymographs without reprocessing the entire dataset, while the “Correlation Analysis” dialog facilitates statistical comparison between kymographs from different fluorescent channels through frame-by-frame Pearson and Spearman correlation analysis with significance testing (Figure 1E). Configuration files can be saved and loaded to maintain consistent analysis parameters across multiple datasets or experimental conditions. The software accepts standard multi-frame TIFF files for both the imaging data and the binary mask (Figure 1A). Multichannel tiff files can be loaded and processed at once (by correctly specifying value in the “Number of Channels” box). All outputs are automatically organized into structured directories within the parent directory of the input images (Figure 1C), ensuring reproducibility and facilitating downstream analysis via built-in Python APIs or custom-written Python/MATLAB programs.

### 2.3 Membrane coordinate extraction

The software begins by loading the user-provided image and binarized mask which can be generated using standard segmentation tools such as Fiji/ImageJ, ilastik, CellProfiler, CellPose, or custom thresholding-based or machine-learning based approaches. For each frame, the software identifies the cell boundary using contour detection algorithm (`skimage.measure.find_contours`) (van der Walt et al., 2014), selects the largest enclosed area to ensure robust handling of imaging artifacts, and computes the geometric centroid. The extracted boundary coordinates are then interpolated to achieve subpixel resolution using piecewise cubic Hermite interpolating polynomial (PCHIP) method (Fritsch and Carlson, 1980), which preserves the shape of the data and prevents oscillations typically observed with higher-order polynomial interpolation. Following interpolation, a circular moving-average filter is applied to smooth the boundary coordinates, reducing high-frequency noise while maintaining the overall boundary morphology. This preprocessing ensures that subsequent alignment and sampling steps operate on geometrically consistent and smooth boundaries across all frames.

### 2.4 Inter-frame boundary alignment

A critical challenge in generating membrane kymographs from time-lapse imaging data is that cell boundaries undergo substantial shape changes and rotations between consecutive frames due to dynamic cellular processes. To address this, the software implements a rotational offset minimization algorithm that aligns boundaries across consecutive frames by exhaustively searching for the optimal angular shift. For the initial frame, the boundary is rotationally aligned to start at a user-specified angular position (default: 180 degrees) by calculating polar angles of all boundary points relative to the centroid and circularly shifting the boundary array to place the desired starting angle at the first index. For subsequent frames, the algorithm determines the best rotational alignment by systematically testing all possible circular shifts of the new boundary relative to the previous frame’s aligned boundary. To handle varying boundary point counts due to cell shape changes, the algorithm subsamples the longer boundary by randomly excluding points until both boundaries contain equal numbers of points, then computes the mean Euclidean distance between corresponding points for each possible shift, and selects the shift that minimizes this distance metric. This approach ensures that homologous membrane regions are consistently tracked across frames despite large-scale morphological changes. The aligned boundaries are stored and used as reference for aligning the next frame, creating a temporally consistent coordinate system throughout the entire time series. This iterative alignment strategy is essential for generating accurate spatiotemporal kymographs, as it prevents artificial discontinuities that would otherwise arise from unaligned boundaries.

### 2.5 Perpendicular intensity sampling

After boundary alignment, the software samples fluorescence intensities in a direction perpendicular to the cell membrane at each boundary point. For each point along the aligned boundary, a local circular arc is fit to a small neighborhood using standard least-squares circle fitting (`circle_fit.standardLSQ`) (Klear and Lauritzen, 2023), which provides a robust estimate of the local membrane curvature and normal direction even in regions of high curvature or irregular boundary geometry. The perpendicular sampling direction is then determined from the vector connecting the fitted circle center to the boundary point. Along this perpendicular direction, fluorescence intensities are sampled at regular intervals (default: 0.5 pixel spacing) extending both inward and outward from the boundary (default: *±*8 pixels, user-adjustable via `l_perp` parameter), using biquadratic interpolation (`scipy.ndimage.map_coordinates` with order=2) (Virtanen et al., 2020) to achieve subpixel accuracy. To reduce sensitivity to single-pixel intensity fluctuations and imaging noise, the software takes the mean of the top 5 intensity values within each perpendicular profile, providing a representative intensity estimate for that membrane position. This sampling process is parallelized across all boundary points using `joblib` with thread-based parallelization, significantly accelerating processing for datasets with long boundaries. The resulting intensity profiles capture both the membrane-localized signal and the immediate cortical region, enabling quantification of proteins and lipids that exhibit enrichment at or near the plasma membrane.

### 2.6 Kymograph generation and normalization

The sampled intensity profiles from all boundary points and all frames are organized into kymograph matrices, where each row represents a specific position along the aligned membrane boundary and each column corresponds to a time point. To reduce temporal noise and reveal underlying intensity dynamics, a locally weighted scatterplot smoothing (LOWESS) algorithm (`statsmodels.nonparametric.smoothers_lowess`) (Cleveland, 1979) is applied along the spatial dimension for each frame, using a smoothing fraction of 0.1 that effectively filters high-frequency spatial variations while preserving genuine membrane features. Following spatial smoothing, the software normalizes each channel’s kymograph using a percentile-based approach: the 2nd and 98th percentiles of the raw (pre-smoothed) intensity distribution across all frames are computed and used to define the normalization range, which effectively excludes extreme outliers while capturing the dynamic range of the signal. The smoothed intensity values are then linearly rescaled to the [0, 1] interval using this percentile range and clipped to prevent values outside this range, ensuring consistent visualization across different channels and experimental conditions. This normalization strategy is particularly effective for live-cell fluorescence data, where photobleaching, temporal variations in illumination, and differences in expression levels can otherwise complicate quantitative comparisons. The normalized kymographs are saved as both `NumPy` files (.npy) for further computational analysis and as Excel spreadsheets (.xlsx) for accessibility to users preferring spreadsheet-based workflows.

### 2.7 Visualization and export

The software generates publication-quality kymograph visualizations with extensive customization options. Users can select from an extensive set of Matplotlib colormaps or use the default Parula colormap (adapted from MATLAB, MathWorks, Natick, MA (MathWorks, 2025)), which provides perceptually uniform color representation suitable for scientific visualization. The intensity scaling can be adjusted interactively through a dedicated GUI panel, allowing users to fine-tune the minimum and maximum intensity values displayed, which is particularly useful for emphasizing specific intensity ranges or revealing subtle spatiotemporal features. The kymographs can be exported in multiple formats: high-resolution raster images (PNG, 300 DPI), vector graphics (SVG and PDF) suitable for publication and further editing in illustration software (such as Inkscape or Adobe Illustrator), and raw data matrices (NumPy.npy and Excel.xlsx formats). Additionally, the software generates preview images showing the tracked boundary and perpendicular sampling lines overlaid on the original fluorescence images, enabling users to visually verify the accuracy of boundary detection, alignment, and intensity sampling. All outputs are organized in clearly structured directories (`Kymo_Processed_Data` and `Kymo_Temporary_Images`) within the same parent directory as the input images, facilitating easy data management and reproducibility.

### 2.8 Multi-channel correlation analysis

For multi-channel datasets, the software includes a built-in correlation analysis module that quantifies the spatiotemporal relationship between different membrane-associated proteins or biosensors. Users can load two kymographs (typically representing different fluorescent channels from the same cell) through an interactive dialog and select correlation parameters. The software computes both Pearson correlation (measuring linear relationships) and Spearman rank correlation (capturing monotonic relationships) between the two kymographs on a frame-by-frame basis. For each time point, the corresponding spatial intensity profiles from both channels are extracted, and optionally, zero-valued pixels can be excluded from the correlation analysis to avoid artifacts from background regions or areas where signal is genuinely absent. Statistical significance is assessed by computing p-values for each correlation coefficient, allowing users to identify time points where the two signals are significantly correlated or anticorrelated. The software generates comprehensive visualizations showing correlation coefficients and p-values as time series plots using `seaborn` for enhanced aesthetics, with shaded confidence bands and significance threshold lines (typically p = 0.05). All correlation results are exported as Excel spreadsheets with columns for frame number, Pearson R, Pearson p-value, Spearman R, and Spearman p-value, enabling further statistical analysis or integration with other analysis pipelines. This correlation analysis capability is particularly valuable for investigating co-localization dynamics, phase relationships, and functional coupling between different signaling components at the cell membrane.

### 2.9 Python API, batch processing, and further analysis

Beyond the GUI-based workflow, the software provides a native Python API (`membrane_kymograph`) that exposes the core processing functionality for programmatic access, enabling integration into custom analysis pipelines and high-throughput batch processing workflows. The API follows a simple object-oriented design: users instantiate a `KymographProcessor` object and call its `process()` method with paths to image and mask files along with desired parameters. The modular architecture allows advanced users to access intermediate processing steps, such as boundary extraction, alignment, and intensity sampling, facilitating custom analyses that build upon the software’s core algorithms.

For batch processing of multiple cells or experimental conditions, users can write Python scripts that iterate over directories of images, apply consistent processing parameters, handle errors gracefully, and aggregate results across multiple datasets. The API supports parallel processing through Python’s multiprocessing capabilities (in addition to `joblib` based built-in multithreading). We provide comprehensive documentation with example scripts demonstrating common batch processing patterns, including processing cells with different parameter requirements (e.g., varying numbers of channels, membrane widths, colormaps, etc. using a csv file), skipping already-processed files to enable interrupted workflow resumption, intensity profiling at specific spatiotemporal locations, and proper error handling. The raw and normalized kymograph data exported as NumPy arrays can be readily imported into scientific Python libraries such as SciPy, scikit-learn, or custom machine learning frameworks for advanced analyses including feature extraction, clustering, dimensionality reduction, and predictive modeling. This programmatic interface significantly extends the utility of the software beyond interactive single-cell analysis, enabling computational researchers to process large-scale datasets and perform sophisticated quantitative analyses that would be impractical through the GUI alone.

### 2.10 Dissecting the dynamics of Ras/PI3K/F-actin network in migrating cells using the software

We have validated the software using microscopy images from cells across phylogenic tree and diverse set of membrane-associated proteins/lipids, in different physiological conditions: Combining GUI and API modes, *>* 100 cells have been processed using this software, as they were migrating or making macropinosomes. As a proof of concept, here we present the data analysis from *Dictyostelium* amoeba, human neutrophils, and murine macrophages which collectively help us to dissect the evolutionarily conserved Ras/PI3K/F-actin network during amoeboid migration, either under stochastic or receptor-driven activation. One key aspect of the migration is the symmetry breaking at the plasma membrane-cortex as specific components asymmetrically localize to front-state/protrusion or back/basal-state regions of the membrane (Banerjee et al., 2025; Ridley et al., 2003). These asymmetric distributions play key role in organizing cellular polarity and regulating migration. Here, our membrane kymographs and correlation analysis show that tumor suppressor protein PTEN and PI(3,4,5)P3 lipid biosensor PH_*Crac*_ consistently maintain tight complementarity as *Dictyostelium* cells migrate (Figure 1D, E; Supplementary Figure 1A). As reported in the previous literature, newly-polymerized F-actin biosensor Lifeact goes through phases of activation and extinction as HL-60 human neutrophil changes shape and migrate (Supplementary Figure 1B, C). Our kymographs demonstrate that when neutrophils go through contractions, Lifeact remained largely absent from the membrane-cortex, whereas during protrusion phases, Lifeact shows strong membrane recruitment (Supplementary Figure 1C). Furthermore, Ras activation biosensor Raf_*RBD*_ and PI(3,4,5)P3 biosensor PH_*Crac*_ show strong positive correlation with slight phase difference (Supplementary Figure 1D-F), indicating that Ras activity positively regulates PI3K activity at the leading edge of migrating cells. While examples so far dissected the dynamics as the Ras/PI3K/F-actin network undergoes stochastic activation, we also studied the receptor driven network activation using RAW 264.7 macrophages. Upon global activation of C5aR receptors in these murine macrophages, PI(3,4,5)P3 biosensor PH_*Akt*_ shows robust membrane recruitment across the entire boundary (Supplementary Figure 1G-I), consistent with the known role of C5aR in activating PI3K signaling. The temporal profiles extracted from the kymographs clearly show a sharp increase in mean membrane-associated PI(3,4,5)P3 levels immediately following C5aR stimulation and eventual adaptation (Supplementary Figure 1I), as expected. All these observations and inferences are consistent with previous findings in the field, validating the accuracy and reliability of our software. Moreover, the kymographs reveal that these spatial patterns are highly dynamic, with rapid transitions between front and back states occurring over timescales of seconds to minutes, highlighting the complex, internal feedback loop mediated regulation of membrane-associated signaling during cell migration. Overall, these analyses demonstrate the utility of *Membrane Kymograph Generator* for quantitatively dissecting the spatiotemporal dynamics of key signaling and cytoskeletal networks at the plasma membrane-cortex in live amoeboid cells.

## 3 DISCUSSION

We have developed *Membrane Kymograph Generator*, a free, open-source, user-friendly, cross-platform software that addresses a significant gap in quantitative live-cell imaging analysis by fully automating the generation and downstream analysis of kymographs along dynamic cell boundaries. This software removes substantial technical barriers that have historically limited membrane dynamics analysis to specialized computational researchers or forced researchers to rely on time-consuming, manual, error-prone approaches. Furthermore, the software’s modular Python architecture, featuring both an accessible GUI for routine use and a comprehensive Python API for programmatic access: this accommodates users ranging from experimentalists seeking straightforward point-and-click kymograph generation and built-in analyses tools to computational researchers requiring high-throughput batch processing and integration with custom analysis pipelines.

At present, *Membrane Kymograph Generator* works as a native desktop software. While this has several advantages, in future, a cloud-native version can be developed to enable analysis of very large datasets that exceed local computational resources and require collaborative sharing. Additionally, the software currently needs a binary mask of only one cell to be present in the field of view, as when more than one disconnected components are present in a masked image, only the component with the largest area is processed and rests are ignored. In future, the software can be tuned to process multiple labelled cells from single binary image mask. In subsequent versions of the software, we can also potentially integrate additional advanced quantitative analysis modules, such as automated detection of spatiotemporal features and their temporal evolution, frequency analysis of membrane oscillations, machine learning-based feature extraction for phenotypic classification, etc.; currently, these or other similar downstream analyses can only be performed with custom-written programs that take advantage of our Python API.

In summary, *Membrane Kymograph Generator* provides the cell biology and biophysics research communities with a much-needed tool for automated, reproducible, and quantitative analysis of membrane dynamics from live-cell imaging data. Its combination of robust algorithms, user-friendly design, cross-platform availability, and open-source licensing makes it accessible to a broad range of researchers and positions it as a valuable addition to the growing ecosystem of open-source bioimage analysis tools. We anticipate that this software will facilitate new discoveries regarding the spatiotemporal organization and regulation of membrane-associated signaling and cytoskeletal processes across diverse cellular systems and physiological contexts.

## ACKNOWLEDGEMENTS

We thank all the members of the Iglesias and Devreotes laboratories (Johns Hopkins University) for their helpful feedback. We especially thank D. Biswas (Iglesias Laboratory, JHU) for his contribution during the development of this software. We would also like to thank Y. Deng, D. S. Pal, and S. Ye (Devreotes Laboratory, JHU) for their help during testing the software. This work was supported by NIH grant nos R35 GM118177 (to P.N.D.) and R01 GM149073 (to P.A.I.). This work is also supported by AFOSR MURI FA95501610052 (to P.N.D.) as well as NIH grant S10 OD016374 (to S. Kuo of the JHU Microscope Facility). This manuscript is the result of funding in whole or in part by the National Institutes of Health (NIH). It is subject to the NIH Public Access Policy. Through acceptance of this federal funding, NIH has been given a right to make this manuscript publicly available in PubMed Central upon the Official Date of Publication, as defined by NIH.

---

**Supplementary Figure 1.**
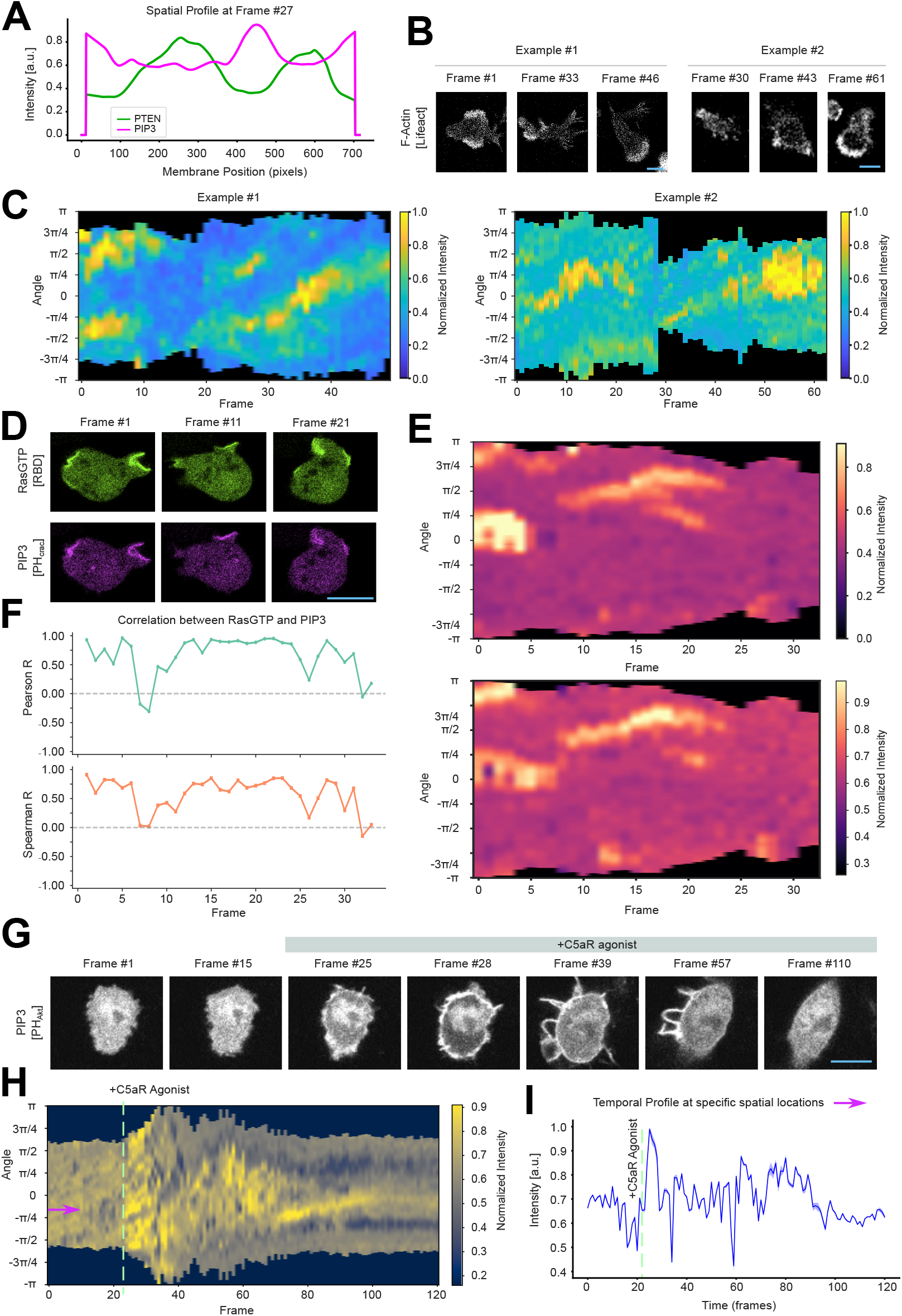
Membrane kymographs and downstream analyses across different cell types using Membrane Kymograph Generator. **(A)** Spatial profile of PTEN (green) and PI(3,4,5)P3 biosensors (magenta) along the membrane boundary of a migrating *Dictyostelium* cell, at a particular frame. The kymographs generated in Figure 1D were used as the data. A similar example has been documented in Python API section of the wiki. **(B, C)** Representative live-cell time-lapse fluorescence images (B) and membrane kymographs (C) of HL-60 human neutrophil cells expressing newly-polymerized F-actin biosensor Lifeact, during random migration. The images were acquired at 7 sec/frame rate. In both examples, “Default” colormaps were used. Here and everywhere else, scale bars indicate 10 *μ*m (unless otherwise specified). **(D-F)** Representative live-cell time-lapse fluorescence images (D), membrane kymographs (E), and time-series correlation analysis plots. The outputs in (D) and (E) were generated by the *“Membrane Kymograph Generator”*, from cells shown in (D). Here, *Dictyostelium* cells were co-expressing biosensors for Ras activation (*RBD*_*Raf*1_-GFP) and PI(3,4,5)P3 (*PH*_*Crac*_-mCherry) and the images were acquired at 7 sec/frame rate. The “magma” colormap was used in the membrane kymographs; note that, intensities were adjusted here after generation of kymographs for better representation (using “Adjust Existing Kymograph” module). Pearson R and Spearman R values were computed using “Correlation Analysis” module. **(G-I)** Representative live-cell time-lapse fluorescence images (G), membrane kymographs (H), and temporal profile plots (I). The outputs in (H) were generated by the *“Membrane Kymograph Generator”*, from cells shown in (G) and “cividis” colormap was used in the kymograph (using “Adjust Existing Kymograph” module). Here, RAW 264.7 murine macrophage cells were expressing PI(3,4,5)P3 biosensor (*PH*_*Akt*_-mCherry) and C5aR receptors were globally activated in the middle of the imaging experiment by adding a saturating dose of C5aR agonists (as shown with vertical lines), leading to robust activation of PI3K in the membrane. The images were acquired at 12 sec/frame rate. The temporal profiles of mean intensities (I) were computed from the kymographs in (H), by loading the raw kymograph data and taking average of 175-185 spatial positions along the membrane boundary at each time point. The selected spatial positions at the beginning of time are marked with magenta arrow (H and I). The data in (I) is mean *±* s.e.m. from all those spatial positions. The (G) and (H) panels show that PI(3,4,5)P3 levels rapidly increase at the membrane following C5aR activation, peaking right after post-stimulation before gradually declining towards baseline levels over the next several frames.

